# TCA cycle entry point, growth variability and amino acid utilization in *Alteromonas macleodii* ATCC 27126

**DOI:** 10.64898/2026.03.04.709670

**Authors:** Waseem Bashir Valiya Kalladi, Daniel Sher

## Abstract

Amino acid catabolism is a vital metabolic process in bacteria, providing energy, carbon and potentially nitrogen as resources, and affecting global cycles of these elements. The ability of a bacterium to catabolize an amino acid is often inferred from the presence of the relevant catabolic pathways in its genome, yet the “gene=function” inference is not straightforward. Here, we use growth assays in 96 well plates on individual amino acids and their combinations to directly measure the ability of a model marine bacterium, *Alteromonas macleodii* ATCC 27126, to utilize these resources for growth. With the exception of aspartate and glutamate, which did not support growth in any of our experiments, ATCC 27126 grew on all other amino acids. However, the probability of growth, together with growth yield and rate, differed depending on the entry point of the catabolic pathway to central carbon metabolism, with robust growth occurring only on amino acids catabolized into pyruvate or acetyl CoA. Growth on combinations of two amino acids revealed reproducible patterns, the clearest being inhibition of growth on other amino acids by asparagine, aspartate and their degradation product, oxaloacetate. Finally, growth was different in test tubes compared with 96 well plates. Our results reveal hidden complexity in amino acid utilization and suggest a “TCA-centric” viewpoint for amino acid utilization, perhaps reflecting the high metabolic flexibility of pyruvate and specific regulatory aspects of the TCA cycle in *Alteromonas*.

## INTRODUCTION

Amino acids are an important resource for bacteria. They are the building blocks for proteins, which constitute a large fraction of living biomass, as well as of other biomolecules. When proteins are released, e.g. during cell death, amino acids become part of the bioavailable organic matter, and can serve as both nitrogen (N) and carbon (C) sources in a wide range of environments^1–3^. In some cases, amino acids can serve as sole carbon and energy sources, particularly when other substrates such as sugars are depleted^4–7^. From an ecological point of view, the catabolism of amino acids provides an important link between organic and inorganic pools of elements such as carbon and nitrogen^8,9^.

Bacteria catabolize amino acids through two major routes: one-carbon metabolism and amino acid–specific catabolic pathways. One-carbon metabolism channels single-carbon units derived from amino acids such as serine, glycine, and histidine into a folate-dependent network that supports nucleotide biosynthesis, methylation reactions, and redox balance via tetrahydrofolate-linked one-carbon carriers^10,11^. In contrast, amino acid–specific catabolic pathways convert individual amino acids through reactions like transamination and oxidative deamination into α-keto acids that are key intermediates in central carbon metabolism (for example pyruvate, acetyl-CoA, 2-oxoglutarate, or succinyl-CoA). Through these pathways, amino acids are converted into molecules that feed central metabolic routes such as the TCA cycle, thereby incorporating dissolved organic matter into cellular energy and biosynthesis^4–6^. Importantly, only amino-acid specific catabolism pathways allow direct acquisition of carbon skeletons from L-amino acids, enabling them to be used as sole carbon sources for growth (one carbon metabolism can be used only to synthesize a limited number of biomass components, or to generate ATP and reducing power)^4,12^. Hence, when growth is observed on an amino acid as a sole carbon source, this is taken as evidence either for the activity of the specific catabolic pathway, or for carbon fixation^13^.

Over past decades, numerous studies have tested bacterial growth on individual L-amino acids across diverse taxa^7,14,15^. A recurring observation is that while some amino acids are readily used as carbon sources, others appear inaccessible to certain bacteria. This suggests that most bacterial species rely only on a subset of amino acids for carbon acquisition even when these compounds dominate the available nutrient pool.

In marine ecosystems, amino acid–based macromolecules such as peptides and proteins represent a large fraction of organic carbon: 13–37% of particulate organic carbon and 3–4% of dissolved organic carbon in oceanic and coastal waters^8,9,16^. Given the availability of amino acids as resources, their catabolic pathways emerge as essential tools for heterotrophic bacteria—particularly particle-associated bacteria, which are frequently studied for their versatile role in organic matter degradation in the oceans^17–19^. In line with this, we previously reported that amino acids support enhanced biomass generation in marine heterotrophic bacteria compared to neutral sugars^20,21^.

*Alteromonas* are a clade of marine heterotrophic bacteria that is widely distributed, highly culturable, and well-studied both in situ and in vitro^22^. *Alteromonas* grow well in multiple media, and can utilize amino acids and peptides as sole carbon sources^20,21^. Nevertheless, genomic analyses reveal that multiple *Alteromonas* species consistently lack catabolic pathways for cysteine, lysine, ornithine, and in some isolates also tryptophan^23^. Moreover, several of the catabolic pathways that are encoded in the genome of the type strain *Alteromonas macleodii* ATCC 27126 (henceforth ATCC 27126), appear incomplete, i.e. they lack homologs of enzymes known from other bacteria to participate in these pathways^23^ (figure 1). We therefore asked which amino acids can be utilized by ATCC 27126 as sole carbon sources, with the assumption that if it can grow on amino acids without full catabolic pathways, this will suggest alternative, uncharacterized degradation pathways. Instead, our results suggest a more complex picture, where growth on an amino acid depends on where its degradation pathway enters central carbon metabolism, on the presence of other amino acids, and on the specific culturing conditions and growth assay used.

**Figure 1:**
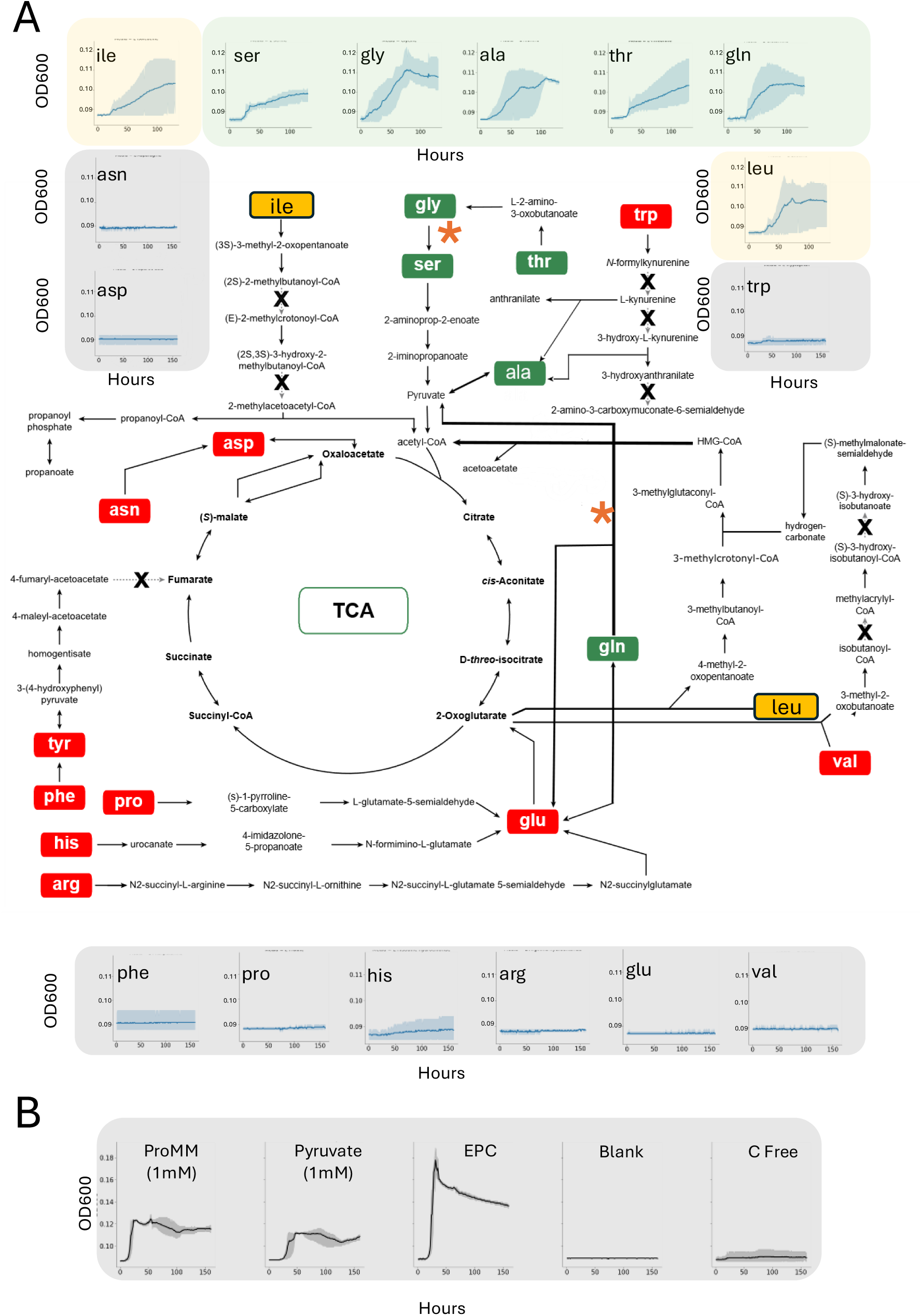
Amino acid catabolism pathways encoded in the genome of Alteromonas macleodii ATCC 2712. A) predicted catabolic pathways, with asterisks showing the reactions identified in the bioinformatic re-annotation. Inset graphs show examples of growth curves on individual amino acids as sole carbon sources, at a concentration of 1mM. Amino acids highlighted in green are considered to be able to support growth, those in yellow, are inconsistent supporters, and those in grey are non-supporters of growth. Note that slow growth was observed on phenylalanine in subsequent experiments, so it is best classified as inconsistent. B) Example of positive and negative control growth curves. EPC – equimolar positive control, i.e. a mixture of all 19 amino acids (0.05 mM each).

## RESULTS AND DISCUSSION

### Refining the bioinformatic predictions of pathways for amino acid degradation in ATCC 27126

Before performing growth experiments on individual amino acids, we first asked whether the genome of ATCC 27126 encodes alternative degradation pathways that circumvent gaps identified in our earlier study (Figure 1A, Supplementary figure 1)^23^. The predicted degradation pathways for seventeen amino acids are shown in Figure 1, with the other three (L-methionine, L-cysteine and L-lysine), not shown as no degradation pathways were found linking them to the TCA cycle. L-methionine was seen to enter the purine ribonucleoside degradation pathway through the “L-methionine degradation I (to L-homocysteine)” pathway, missing direct links to central carbon metabolic pathways, and the other two (L-cysteine and L-lysine) did not have degradation pathways listed under the ATCC 27126 PDGB on BioCyc.

We identified a reaction previously described in *E. coli*^24^ whereby glycine could be metabolized to serine (with a concomitant conversion of tetrahydrofolate to 5,10- methylenetetrahydrofolate) through the activity of the enzymes serine hydroxymethyltransferase (loci RS12410 & RS02225 in the ATCC 27126 genome). Serine is then metabolized in two additional steps to pyruvate (Figure 1A, Supplementary Figure 1). In addition to linking glycine to the tricarboxylic acid (TCA) cycle, this pathway also suggested a potential route for the catabolism of threonine which circumvents the missing reaction converting 2-oxanobutanoate to propionyl-CoA, and the need to assume a way for propionyl-CoA or propanoate to enter core metabolism (Supplementary Figure 1).

Additionally, we identified a metabolic reaction linking glutamine to the TCA cycle through the glutamate dehydrogenase, which produces pyruvate and 2-oxoglutarate from glutamine and chorismate. This reaction, known as “Glutamine degradation I” in BioCyc^25^, comes in addition to the multiple reactions converting glutamine to glutamate, which in turn can then enter the TCA cycle through 2-oxoglutarate. Together, the reannotation of the amino acid degradation pathways suggested plausible pathways for the degradation of two amino acids previously considered to have gaps (Gly, and Thr), and added an alternative reaction for the catabolism of another amino acid, Gln.

### Measurable growth on amino acids in 96 well plates depend on the entry point into core metabolism

We next grew ATCC 27126 on each of 19 individual amino acids as sole carbon sources (L-tyrosine was excluded because of insolubility in water). Relatively low concentrations were used (see below) for several reasons: a) in order to verify that cessation of growth is due to carbon starvation (i.e. the media has a C:N molar ratio of 5:4); b) to avoid precipitation and coloration of amino acids in aqueous solution; c) to avoid potential amino acid toxicity (Supplementary Figure 2). Importantly, comparing growth between different amino acids requires that a standard concentrate on is provided, yet each amino acid is different, comprising different numbers of carbon and nitrogen atoms. Each amino acid also requires a different number of metabolic reactions to reach central carbon metabolism, which could translate into different costs for utilization^26^. We therefore performed five different growth experiments, three with each amino acid presented at the same molar concentration (1mM), and two where the same molar concentration of N atoms was added (0.8mM) (Supplementary Figure 2). Each experiment included also two positive controls – an equimolar mixture of all 19 amino acids (EPC) and 1mM pyruvate, as well as a negative control with no carbon source (Figure 1B, Figure 2).

**Figure 2:**
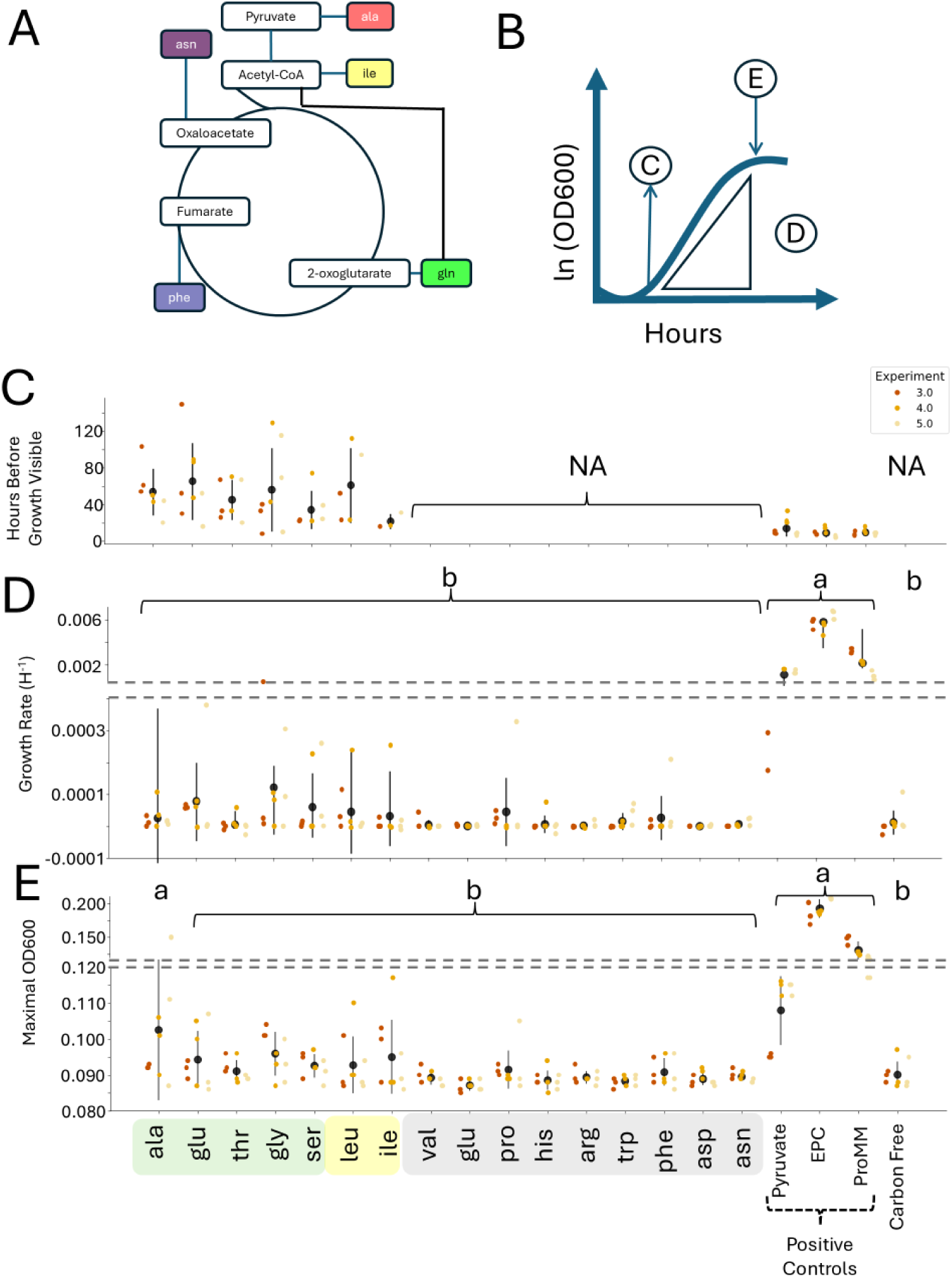
Growth of ATCC 27126 on individual amino acids. A, B) Schematic illustrations of the entry points into the TCA cycle and of the measured parameters (lag phase, growth rate and maximal yield, marked as panels C-E, below). C-E) Different measures of growth on individual amino acids: Lag time (C), growth yield (maximum OD600nm, D) and maximum growth rate (E; are shown for growth on 1mM of each individual amino acid. Results are shown from 3 independent experiments (different shades), each with triplicate wells. NA – no growth observed. Different lowercase letters (a–b) indicate statistically distinct groups based on Kruskal–Wallis and Mann–Whitney U tests (p < 0.05, Bonferroni-corrected); media sharing the same letter do not differ significantly. The amino acids differed significantly from each other in yield (Kruskal-Wallis Test H: 137.215, p: 1.189e-18) and maximum growth rate (Kruskal-Wallis Test H: 96.996, p: 2.151e-11).

Overall, growth differed between the various amino acids in multiple aspects. Firstly, while two amino acids consistently did not show growth (Asp and Glu), for all other amino acids growth was observed in at least one of the individual replicates across all experiments (Figure 1, Supplementary Figure 3). Growth was also observed in 3 out of the 15 replicates of the negative control, consistent with previous studies^20^, possibly due to incomplete drawdown of cellular energy and carbon storage during the 24-hour pre-experiment starvation step. We therefore divided the amino acids into “growth” (growth in >9/15 replicates), “inconsistent” (growth in 7/15 replicates) and “no growth” (<=3/15 replicates in this experiment) (colors in Figure 1). There were also differences between the amino acids in the maximum yield (absorbance at 600nm), growth rate, and lag time (Figure 2B, C).

The patterns of growth on individual amino acids seemed visually to be related to the entry point of each degradation pathway into core metabolism (Figure 1, 2). All of the amino acids that consistently grew were predicted to be catabolized to pyruvate (including Gln, if degraded through the Glutamine degradation I pathway)(Figure 2C). There are two additional amino acids, Trp and Val, that are also predicted to be catabolized to pyruvate yet did not support the growth of ATCC 27126. The predicted catabolism pathways for each of these amino acids contains two gaps (Figure 1). Ile and Leu both inconsistently supported growth, and both are catabolized into acetyl CoA. All other amino acids, which did not support growth, were predicted to be catabolized into metabolites that are downstream from pyruvate and acetyl CoA, namely TCA cycle intermediates such as 2-oxoglutarate, fumarate and oxaloacetate.

Growth on individual amino acids was much slower and less efficient (i.e. reached a lower OD600) compared to an equimolar mixture of all amino acids (Figure 2D, E). It was also slower and less efficient than growth on a mixture of small organic substrates (ProMM media, containing lactate, acetate, pyruvate and glycerol), or on pyruvate alone. Faster growth and higher yield on pyruvate compared to most amino acids that are catabolized to it (with the exception of yield on Ala) suggests a higher metabolic cost for amino acid catabolism compared with growth on the carbon backbone, despite the fact that the amino acids also contain a nitrogen atom that could be used for various anabolic processes (the growth media contained 800μM NH4+, and 50μM PO43-, with a C:N:P ratio of 20:16:1, and hence growth ceased likely due to C starvation). Nevertheless, growth on the carbon backbones only (ProMM and pyruvate) was slower and less efficient than on the equimolar mix of amino acids at the same concentration. This suggests that, when a wider variety of amino acids was present, the increased metabolic cost associated with amino acid degradation was offset either by the ability of the cell to use multiple catabolic pathways in parallel, or to directly incorporate amino acids into growing biomass.

Importantly, while growth on ProMM, pyruvate and the amino acid mixture was almost immediate (typically observed after ∼12 hours), growth on some individual amino acids such as serine, alanine and glutamine could typically be observed only after 24-80 hours (Figure 2C). This was sometimes followed by a burst of relatively fast growth (e.g. Gln in Figure 1), suggesting an extended lag phase where the cells modify their metabolism to deal with the single resource. It also highlights the need for long-term experiments to assess growth, beyond the 24 or 48 hours often used for growth assays with heterotrophic bacteria^27^.

### Combining two amino acids with different entry points into the TCA cycles alters growth

We next asked whether the increased growth on multiple amino acids compared to individual ones could be explained by the mixture having multiple entry points into the TCA cycle. Previous studies have shown that growth increases upon co-feeding with multiple central carbon metabolites in bacteria^7,28^ and yeast^29^. Such co-feeding could, for example, reduce the need for anaplerotic reactions to replenish TCA cycle intermediates used for biosynthesis, and/or reduce the potential for accumulation of some intermediates to toxic levels^30^. We therefore performed growth experiments using equimolar combinations of two amino acids, each representing a different entry point to the TCA cycle (Figure 3A): L-isoleucine, L-asparagine, L-phenylalanine, L-glutamine and L-alanine, entering the TCA cycle through acetyl-CoA, oxaloacetate, fumarate, 2-oxoglutarate, and pyruvate, respectively.

**Figure 3:**
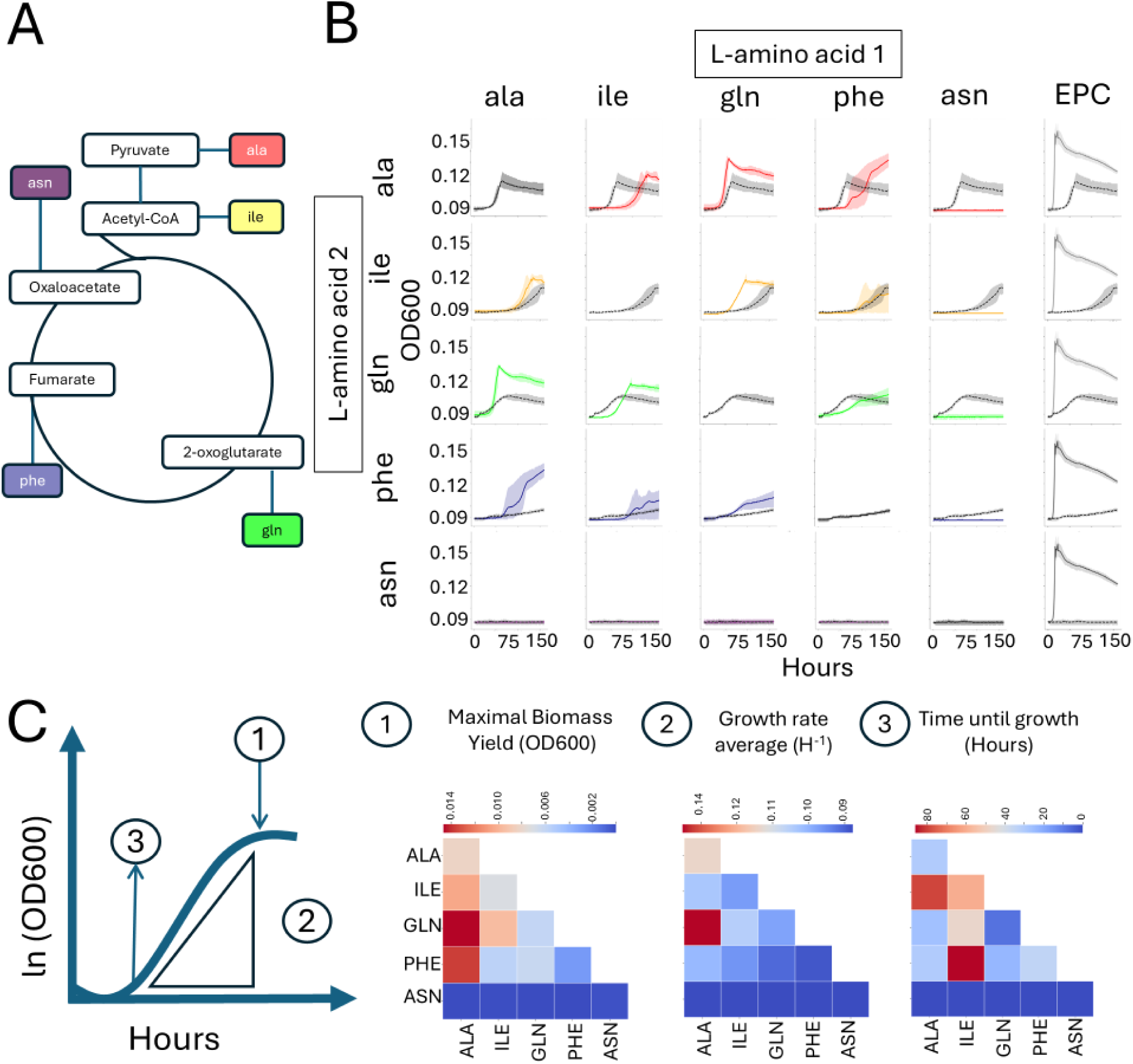
Co-feeding of amino acids. (A) The amino acids, with their corresponding TCA cycle entry points. (B) A matrix of growth curves showing the impact of the addition of amino acid 1 (Columns), on amino acid 2 (Rows). The grey dotted curves in each panel show the growth of amino acid 2 alone (rows, 1mM), and the colored lines show the combined growth of amino acid 2 and 1 (i.e. 0.5mM each of the amino acids in the rows and columns). The grey solid curve in the last row denotes the EPC. (C) Schematic illustration and heatmap of various growth characteristics.

In contrast to our initial hypothesis, we did not observe a simple case of better growth when two amino acids were provided instead of one. Rather, some of the amino acids had qualitatively and quantitatively different effects on growth with others (Figure 3B, C). Alanine and glutamine clearly increased both growth rate and growth yield when added to every amino acid apart from asparagine. Isoleucine and phenylalanine both delayed the growth on most other amino acids (i.e. increased the lag phase), while also increasing the yield on some amino acids but not others (in some cases, particularly when one of the amino acids was phenylalanine, growth continued beyond the ∼165 hours of the experiment, so the final yield was not known). In striking contrast to the other four amino acids, asparagine was unable to support growth by itself and consistently inhibited growth in combination with each of the other four amino acids, with no recovery observed even after 165 hours of monitoring.

When considered from the point of view of the “recipient” amino acid, yield on phenylalanine (which grew very slowly alone in this experiment, and was defined as “not growing” in the experiments described above) always increased when an additional amino acid was added. Alanine and glutamine, which grew robustly alone, each “benefitted” when two amino acids were added, and the results for isoleucine (“inconsistent” when grown alone) were less clear (very minor benefit). There was no clear correlation between the entry point into the TCA cycle and the phenotype of each amino acid, for example, alanine and isoleucine, which both enter the TCA cycle through acetyl CoA, were different as both “affectors” and “affected”. Additionally, the growth on the equimolar combination of all 19 amino acids occurred much earlier than on any combination of two amino acids, growth was faster, and the final yield was higher (Figure 3B). Taken together, these findings indicate that most combinations of two amino acids (each entering the TCA cycle at a different location) are synergistic, but their combined effect still supports much less efficient growth than a mixture of all amino acids, and strongly depends on the specific amino acids and the pairing context.

### Inhibitory effect of asparagine, aspartate and oxaloacetate

Intrigued by the inhibitory effect of asparagine on growth fueled by other amino acids, we asked whether similar inhibition is observed when asparagine is replaced by aspartate or oxaloacetate, the two metabolites downstream in the asparagine catabolism pathway. If so, this could further support the hypothesis that the impact of individual amino acids on growth is determined, at least to some extent, by how their degradation pathways interact with core metabolism. In this experiment, asparagine inhibited, but did not completely abolish, growth on isoleucine, glutamine or phenylalanine (Figure 4). The inhibitory effect increased along the catabolic pathway, with a stronger effect of aspartate and complete inhibition with oxaloacetate.

**Figure 4:**
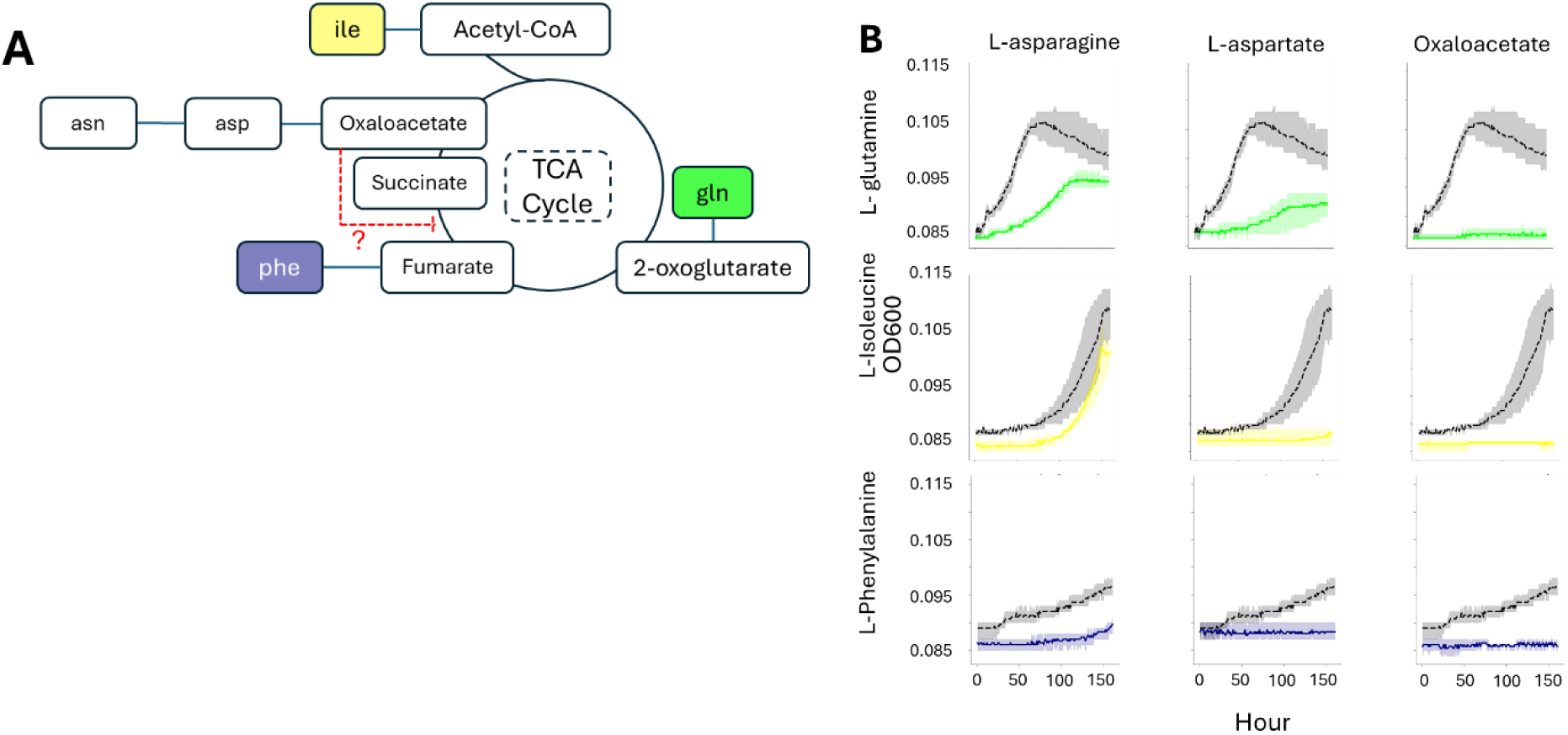
Inhibitory nature of asparagine and downstream metabolites. (A) TCA cycle, showing the entry points of the various amino acids, tested for inhibition by Asparagine and its downstream metabolites. The red inhibition arrow shows the suggested inhibitory effect of oxaloacetate on the succinyl dehydrogenase enzyme. (B) The inhibitory effect of Asparagine derivatives (amino acid 2, columns), against the tested amino acids (amino acid 1, rows). The grey curves in each panel show the growth of amino acid 1 alone (rows, 1mM), and the colored lines show the combined growth of amino acid 2 and 1 (i.e. 0.5mM each of the amino acids in the rows and columns).

The observation of a clear gradient in growth inhibition along the asparagine degradation pathway (asparagine < aspartate < oxaloacetate) supports a model in which metabolic flux toward the downstream TCA cycle underpins the observed growth reduction. This pattern reinforces the concept that the inhibitory effect is not due to external substrate toxicity but arises from intracellular accumulation of key metabolites that disrupt the finely tuned balance of central metabolism. A potential explanation for the specific inhibition by asparagine and its degradation products could be the accumulation of keto-oxaloacetate. This metabolite is the form of oxaloacetate used by the enzyme as part of the TCA cycle, but also spontaneously tautomerizes to enol-oxaloacetate. In *Escherichia coli*, enol-oxaloacetate in turn inhibits succinyl-dehydrogenase, the housekeeping TCA cycle enzyme responsible for the formation of fumarate from succinate^31^.

### Differences in growth on individual amino acids between 96 well plates and test tubes

As a first step towards future experiments that would identify the biochemical and metabolic causes of the differences in growth between amino acids, we scaled up the cultures from 96 well plates (200μl) to borosilicate test tubes (30ml) for 5 individual amino acids, as well as positive and negative controls. We selected two amino acids that revealed robust growth in 96 well plates (alanine and glutamine), two that were inconsistent (isoleucine, which also has two missing reactions in its predicted catabolism pathway, and asparagine), and one, aspartate, that revealed a strong inhibitory activity. In addition to growth (change in OD600nm, measured in parallel in 96 well plates and test tubes), we also counted cells using two orthogonal methods, flow cytometry (FCM) and plating (colony forming units, CFU), and measured biofilm formation on the test tubes using crystal violet ^21^ (Table 1, Figure 5).

**Figure 5:**
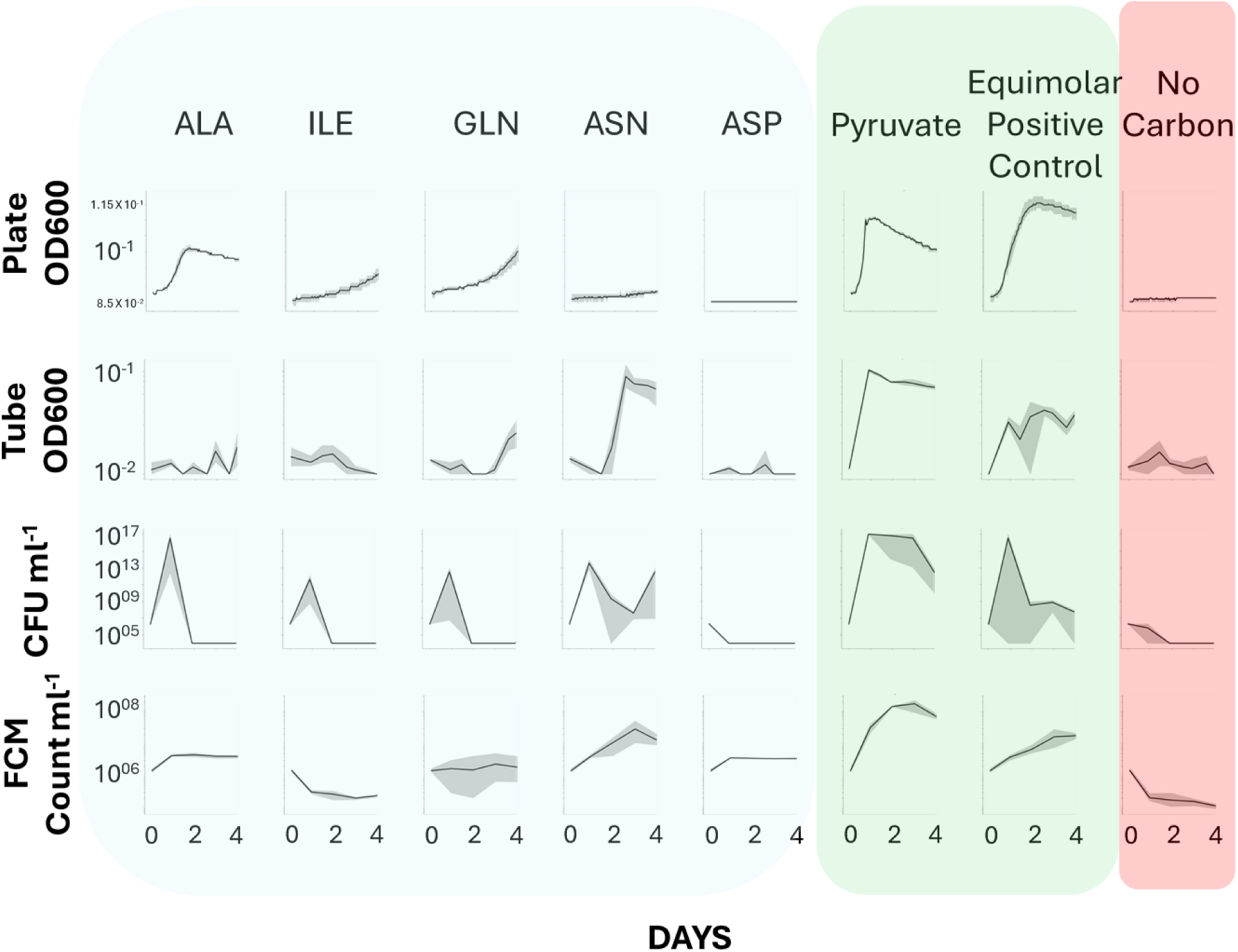
Growth assays in test tubes compared to 96 well plates. From left to right: The growth (OD and cell count) of ATCC 27126 on various amino acids (blue background), in comparison with the pyruvate positive control and the equimolar positive control (green background), and the carbon free negative control (red background). The top row shows growth on plates, in comparison with the OD and cell counts in tubes (Lower three rows).

**Table 1:**
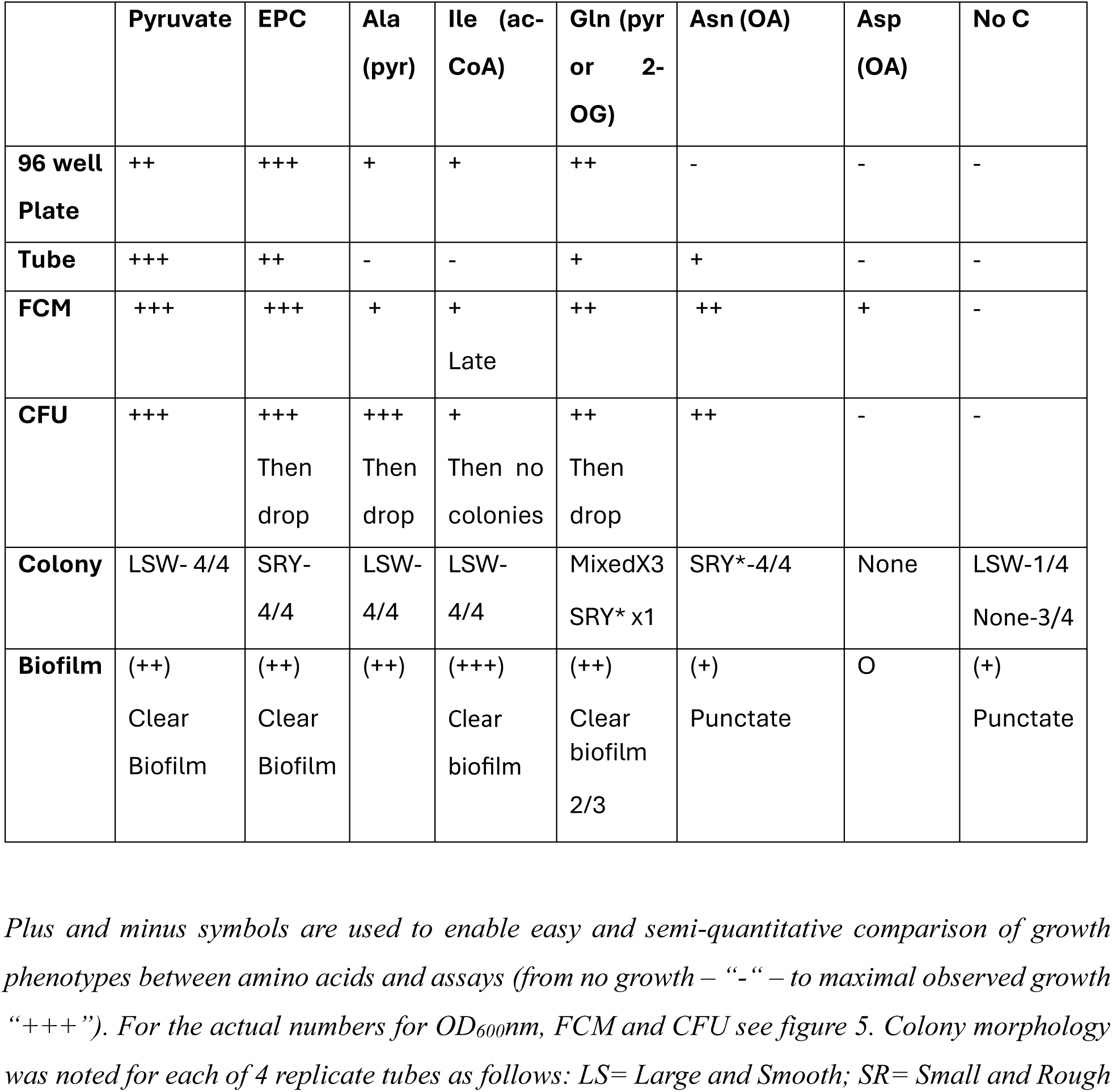
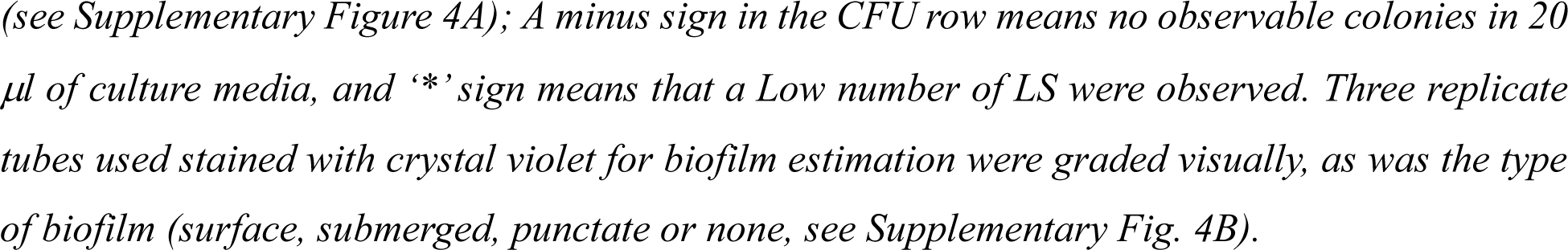
Growth phenotypes in 96 well plates and test tubes.

Growth on the equimolar positive control and on pyruvate was similar between the test tubes and the 96 well plates (Figure 5). Additionally, two amino acids also revealed similar growth phenotypes in the 96 well plates and the tubes – glutamine, which grew after 2-3 days in both conditions, and aspartate, which did not grow under either condition. However, three other amino acids revealed different growth patterns: alanine and isoleucine supported observable growth (increase in OD600nm) only in 96 well plates and not in test tubes, whereas asparagine grew slowly in the 96 well plates, consistent with some of the experiments described above, but rapidly and robustly in the test tubes.

FCM-based cell counts mostly agreed with the changes in OD600nm, increasing in cultures grown on glutamine and asparagine, but less so on alanine, isoleucine and aspartate. However, colony forming units (CFUs) revealed different dynamics, increasing in 3 out of the 5 amino acids after 1 day, but then decreasing to very low numbers. Only asparagine supported increased CFU counts across the 4-day experiment. No colonies were observed from cultures incubated with aspartate, consistent with a strong inhibitory effect.

Intriguingly, we observed two distinct colony morphologies during CFU enumeration: a Large Smooth White (LSW) phenotype and a Small Rough Yellow (SRY) phenotype (Table 1, Supplementary Figure 4A). These phenotypes were different between the different growth substrates, with LSW colonies observed on pyruvate, alanine (which is catabolized to pyruvate) and isoleucine (which is catabolized to acetyl CoA). In contrast, SRY colonies were observed in cultures grown on mixed amino acids and aspartate (which is catabolized to oxaloacetate). Cultures grown on glutamine, which as suggested above can be catabolized to either 2-oxoglutarate or pyruvate, contained both colony morphologies in 3 out of 4 biological replicates. Notably, we used the same marine agar plates for the CFU assays, with colonies observed after 2-3 days. Therefore, the different colony morphology observed in these plates represents a “memory” of the growth conditions the cells experienced in liquid media, which was maintained over at least ∼10-20 generations after they were sowed on the same marine agar plates.

CFU counts consistently exceeded FCM measurements across all tested conditions, a phenomenon potentially attributed to cell aggregation or stickiness characteristic of Gammaproteobacteria, which impairs accurate serial dilution during CFU enumeration^21^. Indeed, floating biofilms have previously been described in *Alteromonas* ^32,33^. In this study we did not measure floating biofilms, but note that in most cultures, a biofilm formed on the sides of the test tubes, as well as (potentially) at the air-surface interface. However, the intensity of the biofilm differed between the cultures grown on different amino acids, as did its appearance (at the surface, as submerged rings, or as “punctate” foci, Table 1, Supplementary Figure 4B). Growth on pyruvate, mixed amino acids, alanine, glutamine and isoleucine resulted in a clear and relatively dense biofilm, whereas on asparagine and in the negative control (no added carbon sources) a weaker, punctate, biofilm was observed. Aspartate cultures failed to form biofilms altogether, consistent with the inhibitory effect of this amino acid on growth.

## DISCUSSION

The question of “which organisms use what resources, and when”, is fundamental to our understanding of microbial life from single cells to ecosystems. The presence of genes encoding the catabolic pathway for a specific resource in the genome of an organism provides evidence that this organism can utilize this resource, and their expression suggests that pathway is active. However, the “gene=function” inference is not straightforward: there are bioinformatic challenges in determining whether a pathway is present and complete (e.g. the gaps in some amino acid catabolism pathways shown in Figure 1^23^), gene expression does not always result in active proteins and pathways^34,35^, and the complexity of metabolic networks means that resource uptake and utilization do not always translate into organismal growth^36^. Therefore, specific assays measuring different aspects of resource utilization are necessary in order to define the “metabolic niche” of an organism. Growth assays, such as those described here, are one approach to directly measure the ability of an organism to utilize a resource for growth, and are often part of the methodology used to describe new bacterial species^37^.

Our results describing the growth of *A. macleodii* ATCC 27126 in 96 well plates on individual amino acids and (some of) their combinations suggest that their utilization for growth may be related to the entry point of their catabolic pathways into core metabolism (the TCA cycle). While this provides a clear “organizing principle” for this aspect of metabolism, we also describe hidden complexity in amino acid utilization – a high level of variability between and within experiments, unexpected interactions between amino acids, and differences in what constitutes “growth” under different experimental conditions.

### Variation between and within experiments is best described as a growth probability

With the exception of two amino acids, aspartate and glutamate, which did not support growth in any of our experiments, ATCC 27126 grew on all other amino acids in at least three of 15 different wells across five different experiments (Supplementary Figure 3). This variability occurred despite our effort to use a simple and reproducible experimental plan, without the use of potentially ambiguous terms (e.g. “mid exponential” or “late exponential” growth), and with an extended starvation period before the start of the experiment aimed to minimize resource carryover. The variability was both between replicate wells from the same experiment and between independent experiments, further supporting the idea that it does not originate from subtle differences in the experimental setup, but rather represents a bona-fide biological phenomenon. The sources of growth variability have been explored to some extent at the single-cell level^38^, but their effect on bulk culture growth, cell physiology, metabolism or gene expression have rarely been studied in detail^39,40^. In experiments where many organisms or conditions are tested, variability is often addressed by selecting the majority of outcomes among experimental replicates^41,42^, accepting a single growing culture as evidence of the ability to grow^26^, or using statistical methods to select only “reproducible” growth curves for further analysis^43^.

Our results suggest that the probability of growth on an individual amino acid is a measurable trait in *A. macleodii* ATCC 27126. This probability, together with growth yield and rate, differs depending on the entry point of the catabolic pathway to central carbon metabolism - growth was robust only on amino acids catabolized into pyruvate or acetyl CoA. The “TCA-centric” viewpoint suggests that regulation of flux through central carbon metabolism, rather than through the individual pathways for catabolizing each amino acid, defines the probability that ATCC 27126 will succeed in utilizing amino acids as single carbon sources. This viewpoint is supported to some extent by the observation that all three metabolites linking asparagine to the TCA cycle (Asn, Asp and Oxaloacetate) inhibit growth on other amino acids.

We hypothesize that the different probabilities for growth on individual amino acids are due to two non-mutually exclusive reasons: the metabolic flexibility of pyruvate, and specific regulatory aspects of the TCA cycle in *Alteromonas*. The high metabolic flexibility of pyruvate, which is at the core of carbon metabolism, has long been recognized^44,45^: it can be used to generate ATP and reducing power through both the TCA cycle and through fermentation to acetate; to replenish TCA cycle intermediates, enabling them to be used for biosynthesis of key biomass components such as amino acids (anaplerosis); and as the initial substrate for gluconeogenesis, producing storage compounds as well as the backbones for nucleotides and cell wall components^46^. The genome of ATCC 27126 encodes all of these potential pathways. We speculate that this flexibility allows cells to more easily adapt to using pyruvate-generating amino acids as sole carbon sources after a period of carbon starvation, compared to amino acids that are catabolized to other TCA cycle intermediates. This increases the probability that the population recovers and starts growing, as well as the rate and yield of subsequent growth.

An alternative hypothesis is that Alteromonas are more sensitive to perturbations in the relative abundance of different TCA cycle intermediates compared with other bacteria. This hypothesis is supported by the initial description of 21 strains of *A. macleodii* from 1972, none of which could grow on any of the TCA cycle intermediates tested (citrate, 2 oxoglutarate, succinate or fumarate, as well as glycolate which is part of a TCA “bypass”)^37^. This is in contrast to many other studied bacteria, which grow on these molecules as sole C sources ^47–49^. Inability to grow on TCA cycle intermediates is usually associated with a partial cycle (e.g. due to gene loss or mutation^50^), although it has also been proposed that in some bacteria increased concentrations of TCA intermediates can lead to dysregulation of the cycle^51,52^. *Alteromonas* encode the full cycle, and therefore it is more likely that their unusual inability to grow on TCA cycle intermediates as sole carbon sources is due to some yet-unknown aspect of the regulation of core metabolism rather than a true inability to utilize these resources.

### Condition-dependent growth in Alteromonas

Growth in bacteria is ostensibly simple to measure – the cultures become turbid, measurable as changes in optical density at 600nm. In many cases, turbidity is used at a single time-point to assess whether growth occurs or not^53,54^. Our results show, firstly, that a lag time of several days can occur even in marine copiotrophic bacteria such as Alteromonas, that grow relatively rapidly. Secondly, some amino acids supported growth differently in 96 well plates and tubes, highlighting how physical culture conditions shape the realized metabolic niche of ATCC 27126.

In 96-well plates, cultures were incubated at small volume at mostly static conditions (except for one out of five replicates where the plates were shaken for 3 seconds before every reading). This enabled us to quantify growth at hourly resolution, resulting in the observed differences in lag time, rate and yield that mapped in part onto the predicted entry point of each catabolic pathway into central metabolism. When the same substrates were scaled up to test tubes, several phenotypes changed qualitatively. Alanine and isoleucine no longer produced a clear increase in OD600, whereas asparagine now supported growth. Cell count methods supplementing optical density showed that this was not a simple loss or gain of viability: flow cytometry and CFU counts increased, and extensive biofilm formation was observed for most substrates on tube walls and, potentially, at the air–liquid interface. One possibility is that that increased culture volume (tube length) and/or altered geometry generate oxygen and nutrient gradients that persist despite tube shaking, and that these gradients favor surface-associated growth and spatially structured populations^55–59^. We propose that under such conditions, biomass can accumulate primarily in biofilms or aggregates that contribute little to bulk OD, decoupling optical turbidity from total cell numbers and masking growth on some amino acids. Nevertheless, we note that no clear difference was observed between plates that were maintained as static (possibly resulting in oxygen depletion) and plates that were shaken every 60 minutes. Therefore, it is unlikely that oxygen concentration alone is responsible for the changes in growth phenotypes between the two conditions.

In addition to potentially affecting the balance between growth in liquid and growth in biofilm, different amino acids also induced different colony morphologies (Large Smooth White and Small Rough Yellow). Strikingly, these phenotypes were maintained despite subsequent growth of the plated bacteria on under uniform conditions (marine agar plates), and therefore reflect a “long term memory” of the previous growth conditions or available resources. It has been reported in other bacteria that alterations in central metabolism and TCA intermediates feed into global regulatory systems such as GMP cycle^51^, which in turn control EPS production^60^, and colony architecture^61^. Similarly, biofilm formation can also change according to the carbon source supplied^57,58^. However, we are unaware of a study that shows such changes persisting over multiple generations (i.e. 2-3 days required to form colonies in our experiment), and thus representing a memory of previous conditions. Further studies are needed to determine whether the changes in colony morphology are the result of a “developmental” program due to changes in gene expression (e.g. conceptually similar to endospore formation^62,63^), or are due to epigenetic changes.

Bacteria in the ocean are unlikely to encounter a single amino acid at millimolar concentration as their sole carbon and energy source; instead, they experience complex, heterogeneous mixtures of dissolved and particulate substrates, often at micromolar or lower levels, and in dynamic microscale gradients. Thus, single amino acid assays emphasize metabolic potential and regulatory constraints under highly simplified conditions, rather than directly mimicking natural resource landscapes. Yet, they highlight a potential “organizing principle” in Alteromonas metabolism, namely the importance of the entry point into the TCA cycle in determining overall metabolism and growth. Additionally, elucidating the specific factors regulating growth under free-living (planktonic) and particle-attached (biofilm) in Alteromonas could be of significant ecological interest: these organisms are usually found attached to particles in nature, where they often “bloom” to form a large fraction of the population^64^, and where their biofilms can modify processes such as particle sinking and carbon export^65^.

## CONCLUSION

This study demonstrates that amino acid catabolism in ATCC 27126 is tightly constrained by the position at which degradation products enter the TCA cycle, with growth observed mainly on amino acids whose carbon is funneled into pyruvate or acetyl-CoA. Amino acids catabolized to downstream TCA intermediates rarely supported consistent growth, highlighting a TCA-centric organization of carbon flow in this bacterium. Co-feeding experiments with selected amino acid pairs further revealed strong, reproducible interactions, including inhibition associated with asparagine, aspartate and oxaloacetate, emphasizing that resource combinations, not only individual substrates, shape growth outcomes. Scale-up experiments in test tubes additionally uncovered two stable colony phenotypes, whose occurrence depended on the prior amino acid substrate and persisted on uniform marine agar, suggesting long-term, resource-imprinted cellular metabolic states. Together, these results highlight hidden complexity in amino acid utilization, suggest preferences for routing carbon through specific TCA entry points, and underscore the need for long-term, fine-scale growth assays that also track phenotypic heterogeneity when linking genomic potential to realized metabolic function in marine heterotrophs.

## MATERIALS AND METHODS

### Media Preparation

Media were prepared fresh for every experiment from pre prepared stock solutions. Recipe described in Supplementary Table was strictly followed. All stocks were filtered sterilized. pH was maintained at neutral. 96 well plates without Amino acids added were pre-incubated for quality control.

### Cell culture and starvation

Alteromonas macleodii 27126 cells were revived from −80°C freezer in 2ml of Marine Broth (BD Difco™ Dehydrated Culture Media: Marine Broth 2216). Cells grown for 36 hours were inoculated to ProMM Media at 1:100 dilution for 200ul volume. Serially, 200ul of Carbon Free ProMM was inoculated from the ProMM culture in the same dilution after 12 hours of growth, in order to starve it off from any carbon reserves for 12 hours. Subsequently this starved culture was used to inoculate all test wells in the various experiments. In the case of the scale up experiment, the dilutions were maintained, but a total volume of 50ml of starved culture was attained. Scale up experiment was set up to attain 30ml of working media volume in the tubes.

### Experimental set up

Plate reader and incubator were set to a temperature of 26°C. 96 well plates were read for OD600 in the plate reader to get biomass proxy. Alternatively, tubes in the scale up experiment were read using a UV-Vis spectroscope compatible with glass culture tubes without transferring to cuvettes. Scale up CFU was done by spot plating 20ul on Marine Agar (BD Difco™ Dehydrated Culture Media: Marine Agar 2216). Scale up samples were collected for FCM at a volume of 100ul every day from each tube aseptic conditions inside a biosafety cabinet. This sample is diluted in a cryo-vial at 10^-1^ in Artificial Sea Water, fixed with glutaraldehyde (2.5%), and subsequently stored in −80°C

### Sensitivity checks and statistics

For the carbon growth curves in figure 1, marking of the start of the growth phase (time until growth), was done manually by reviewing data points and the curves plotted. Double blind marking was done to check for individual bias. A 20-hour window was used to calculate growth rate. Growth rates and maximal yield across amino acid treatments were compared statistically. A Kruskal–Wallis test assessed overall differences, followed by Dunn–Mann–Whitney U post-hoc tests with Bonferroni correction. Mean-based letter groupings were generated to visualize relative similarities among media. Analyses used NumPy, pandas, SciPy, and Pingouin (Code available on Sher Lab GitHub).

### Toxicity Test

Amino acid solutions were prepared at 2.5%, 0.25%, and 0.025% (w/v), and added on top of ProMM, which already contains Pyruvate, glycerol, acetate, and lactate at 0.5% (w/v) each. ProMM without amino acids, served as Positive control and Carbon free ProMM was the negative control. OD was measured at time, 0 hours and 24 hours.

## Acknowledgements

This study was supported by a grant from the National Science Foundation - United States-Israel Binational Science Foundation (NSF-BSF 2246707/2022691) and the Israel Science foundation (grants 1786/20 and 1887/25).

## REFERENCES

1. Morono, Y. et al. Carbon and nitrogen assimilation in deep subseafloor microbial cells. Proc. Natl. Acad. Sci. 108, 18295–18300 (2011).

2. Wang, X., Xia, K., Yang, X. & Tang, C. Growth strategy of microbes on mixed carbon sources. Nat. Commun. 10, 1279 (2019).

3. Bryson, S. et al. Proteomic Stable Isotope Probing Reveals Taxonomically Distinct Patterns in Amino Acid Assimilation by Coastal Marine Bacterioplankton. mSystems 1, e00027–15 (2016).

4. Díaz-Pérez, A. L., Díaz-Pérez, C. & Campos-García, J. Bacterial l-leucine catabolism as a source of secondary metabolites. Rev. Environ. Sci. Biotechnol. 15, 1–29 (2016).

5. Hou, S. et al. Genome sequence of the deep-sea -proteobacterium Idiomarina loihiensis reveals amino acid fermentation as a source of carbon and energy.

6. Wünsch, D. et al. Amino Acid and Sugar Catabolism in the Marine Bacterium Phaeobacter inhibens DSM 17395 from an Energetic Viewpoint. Appl. Environ. Microbiol. 85, e02095–19 (2019).

7. Perrin, E. et al. Diauxie and co-utilization of carbon sources can coexist during bacterial growth in nutritionally complex environments. Nat. Commun. 11, 3135 (2020).

8. Sipler, R. E. & Bronk, D. A. Dynamics of Dissolved Organic Nitrogen. in Biogeochemistry of Marine Dissolved Organic Matter 127–232 (Elsevier, 2015). doi:10.1016/B978-0-12-405940-5.00004-2.

9. Hansell, D. & Carlson, C. Biogeochemistry of Marine Dissolved Organic Matter. (Elsevier, 2024). doi:10.1016/C2022-0-02767-7.

10. Stover, P. J. One-Carbon Metabolism–Genome Interactions in Folate-Associated Pathologies,. J. Nutr. 139, 2402–2405 (2009).

11. Ducker, G. S. & Rabinowitz, J. D. One-Carbon Metabolism in Health and Disease. Cell Metab. 25, 27–42 (2017).

12. Sun, J. et al. One Carbon Metabolism in SAR11 Pelagic Marine Bacteria. PLoS ONE 6, e23973 (2011).

13. Tang, K.-H., Tang, Y. J. & Blankenship, R. E. Carbon Metabolic Pathways in Phototrophic Bacteria and Their Broader Evolutionary Implications. Front. Microbiol. 2, (2011).

14. Ganesan, A. et al. Cloacibacillus evryensis gen. nov., sp. nov., a novel asaccharolytic, mesophilic, amino-acid-degrading bacterium within the phylum ‘Synergistetes’, isolated from an anaerobic sludge digester. Int. J. Syst. Evol. Microbiol. 58, 2003–2012 (2008).

15. Scheifinger, C., Russell, N. & Chalupa, W. Degradation of Amino Acids by Pure Cultures of Rumen Bacteria. J. Anim. Sci. 43, 821–827 (1976).

16. Bianchi, T. S. & Canuel, E. A. Proteins: Amino Acids and Amines. in Chemical Biomarkers in Aquatic Ecosystems.

17. Nguyen, T. T. H. et al. Microbes contribute to setting the ocean carbon flux by altering the fate of sinking particulates. Nat. Commun. 13, 1657 (2022).

18. Boeuf, D. et al. Biological composition and microbial dynamics of sinking particulate organic matter at abyssal depths in the oligotrophic open ocean. Proc. Natl. Acad. Sci. 116, 11824–11832 (2019).

19. Enke, T. N., Leventhal, G. E., Metzger, M., Saavedra, J. T. & Cordero, O. X. Microscale ecology regulates particulate organic matter turnover in model marine microbial communities. Nat. Commun. 9, 2743 (2018).

20. Forchielli, E., Sher, D. & Segrè, D. Metabolic Phenotyping of Marine Heterotrophs on Refactored Media Reveals Diverse Metabolic Adaptations and Lifestyle Strategies. mSystems 7, e00070–22 (2022).

21. Valiya Kalladi, W. B., et al. Biological insights and methodological challenges learned from working with a diverse heterotrophic marine bacterial library. Preprint at 10.1101/2025.11.25.689180 (2025).

22. Wietz, M., López-Pérez, M., Sher, D., Biller, S. J. & Rodriguez-Valera, F. Microbe Profile: Alteromonas macleodii − a widespread, fast-responding, ‘interactive’ marine bacterium. Microbiology 168, (2022).

23. Sher, D. et al. Collaborative metabolic curation of an emerging model marine bacterium, Alteromonas macleodii ATCC 27126. PLOS One 20, e0321141 (2025).

24. Stauffer, G. V. Regulation of Serine, Glycine, and One-Carbon Biosynthesis. EcoSal Plus 1, 10.1128/ecosalplus.3.6.1.2 (2004).

25. Karp, P. D. et al. The BioCyc collection of microbial genomes and metabolic pathways. Brief. Bioinform. 20, 1085–1093 (2019).

26. Flickinger, S. F. & Gralka, M. Biosynthesis cost and catabolic pathway length explain frequency of amino acid utilization as carbon sources by bacteria. Preprint at 10.1101/2025.10.25.684513 (2025).

27. Clerc, E. E. et al. Chemotaxis, growth, and inter-species interactions shape early bacterial community assembly. ISME J. 19, wraf101 (2025).

28. Park, J. O. et al. Synergistic substrate cofeeding stimulates reductive metabolism. Nat. Metab. 1, 643–651 (2019).

29. Li, X. et al. Modular deregulation of central carbon metabolism for efficient xylose utilization in Saccharomyces cerevisiae. Nat. Commun. 16, 4551 (2025).

30. Krüger, A. et al. Impact of CO_2_ /HC_3_^−^ Availability on Anaplerotic Flux in Pyruvate Dehydrogenase Complex-Deficient Corynebacterium glutamicum Strains. J. Bacteriol. 201, (2019).

31. Zmuda, A. J. et al. A universal metabolite repair enzyme removes a strong inhibitor of the TCA cycle. Nat. Commun. 15, 846 (2024).

32. Robertson, J. M. et al. Marine bacteria *Alteromonas* spp. require UDP-glucose-4-epimerase for aggregation and production of sticky exopolymer. mBio 15, e00038–24 (2024).

33. Givati, S. et al. Diversity in the utilization of different molecular classes of dissolved organic matter by heterotrophic marine bacteria. Appl. Environ. Microbiol. 90, e00256–24 (2024).

34. Wettstadt, S. & Llamas, M. A. Role of Regulated Proteolysis in the Communication of Bacteria With the Environment. Front. Mol. Biosci. 7, 586497 (2020).

35. Sinha, A. K., Laursen, M. F. & Licht, T. R. Regulation of microbial gene expression: the key to understanding our gut microbiome. Trends Microbiol. 33, 397–407 (2025).

36. Takano, S., Vila, J. C. C., Miyazaki, R., Sánchez, Á. & Bajić, D. The Architecture of Metabolic Networks Constrains the Evolution of Microbial Resource Hierarchies. Mol. Biol. Evol. 40, msad187 (2023).

37. Baumann, L., Baumann, P., Mandel, M. & Allen, R. D. Taxonomy of Aerobic Marine Eubacteria. J. Bacteriol. 110, 402–429 (1972).

38. Lindemann, D., Westerwalbesloh, C., Kohlheyer, D., Grünberger, A. & Von Lieres, E. Microbial single-cell growth response at defined carbon limiting conditions. RSC Adv. 9, 14040–14050 (2019).

39. Biller, S. J., Coe, A., Roggensack, S. E. & Chisholm, S. W. Heterotroph Interactions Alter *Prochlorococcus* Transcriptome Dynamics during Extended Periods of Darkness. mSystems 3, e00040–18 (2018).

40. Roller, B. R. K. et al. Single-cell mass distributions reveal simple rules for achieving steady-state growth. mBio 14, e01585–23 (2023).

41. Long, R. A. & Azam, F. Antagonistic Interactions among Marine Pelagic Bacteria. Appl. Environ. Microbiol. 67, 4975–4983 (2001).

42. Cordero, O. X. et al. Ecological Populations of Bacteria Act as Socially Cohesive Units of Antibiotic Production and Resistance. Science 337, 1228–1231 (2012).

43. Sher, D., Thompson, J. W., Kashtan, N., Croal, L. & Chisholm, S. W. Response of *Prochlorococcus* ecotypes to co-culture with diverse marine bacteria. ISME J. 5, 1125–1132 (2011).

44. Echlin, H., Frank, M., Rock, C. & Rosch, J. W. Role of the pyruvate metabolic network on carbohydrate metabolism and virulence in *Streptococcus pneumoniae*. Mol. Microbiol. 114, 536–552 (2020).

45. Zhang, Y. et al. Model-guided metabolic rewiring to bypass pyruvate oxidation for pyruvate derivative synthesis by minimizing carbon loss. mSystems 9, e00839–23 (2024).

46. Dağ, Ç. & Kahraman, K. Exploring the biochemical landscape of bacterial medium with pyruvate as the exclusive carbon source for NMR studies. J. Biomol. NMR 79, 143–153 (2025).

47. Liu, M. et al. CitAB Two-Component System-Regulated Citrate Utilization Contributes to *Vibrio cholerae* Competitiveness with the Gut Microbiota. Infect. Immun. 87, e00746–18 (2019).

48. Pos, K. M., Dimroth, P. & Bott, M. The *Escherichia coli* Citrate Carrier CitT: a Member of a Novel Eubacterial Transporter Family Related to the 2-Oxoglutarate/Malate Translocator from Spinach Chloroplasts. J. Bacteriol. 180, 4160–4165 (1998).

49. Warner, J. B. & Lolkema, J. S. Growth of Bacillus subtilis on citrate and isocitrate is supported by the Mg2+–citrate transporter CitM. Microbiology 148, 3405–3412 (2002).

50. Zhou, H. et al. A citric acid cycle-deficient Escherichia coli as an efficient chassis for aerobic fermentations. Nat. Commun. 15, 2372 (2024).

51. Yuan, X. et al. Tricarboxylic Acid (TCA) Cycle Enzymes and Intermediates Modulate Intracellular Cyclic di-GMP Levels and the Production of Plant Cell Wall–Degrading Enzymes in Soft Rot Pathogen *Dickeya dadantii*. Mol. Plant-Microbe Interactions® 33, 296–307 (2020).

52. Chittezham Thomas, V., et al. A Dysfunctional Tricarboxylic Acid Cycle Enhances Fitness of Staphylococcus epidermidis During β-Lactam Stress. mBio 4, e00437–13 (2013).

53. Thompson, L. J. et al. Gene Expression Profiling of *Helicobacter pylori* Reveals a Growth-Phase-Dependent Switch in Virulence Gene Expression. Infect. Immun. 71, 2643–2655 (2003).

54. Smakman, F. & Hall, A. R. Exposure to lysed bacteria can promote or inhibit growth of neighboring live bacteria depending on local abiotic conditions. FEMS Microbiol. Ecol. 98, fiac011 (2022).

55. Depetris, A. et al. Morphogenesis and oxygen dynamics in phototrophic biofilms growing across a gradient of hydraulic conditions. iScience 24, 102067 (2021).

56. Chang, Y.-W. et al. Biofilm formation in geometries with different surface curvature and oxygen availability. New J. Phys. 17, 033017 (2015).

57. Klausen, M. et al. Biofilm formation by *Pseudomonas aeruginosa* wild type, flagella and type IV pili mutants. Mol. Microbiol. 48, 1511–1524 (2003).

58. Sauer, K. et al. The biofilm life cycle: expanding the conceptual model of biofilm formation. Nat. Rev. Microbiol. 20, 608–620 (2022).

59. Oliveira, N. M. et al. Biofilm Formation As a Response to Ecological Competition. PLOS Biol. 13, e1002191 (2015).

60. Poulin, M. B. & Kuperman, L. L. Regulation of Biofilm Exopolysaccharide Production by Cyclic Di-Guanosine Monophosphate. Front. Microbiol. 12, 730980 (2021).

61. Ojha, A. K., Mukherjee, T. K. & Chatterji, D. High Intracellular Level of Guanosine Tetraphosphate in *Mycobacterium smegmatis* Changes the Morphology of the Bacterium. Infect. Immun. 68, 4084–4091 (2000).

62. Veening, J.-W. et al. Bet-hedging and epigenetic inheritance in bacterial cell development. Proc. Natl. Acad. Sci. 105, 4393–4398 (2008).

63. Veening, J.-W., Smits, W. K. & Kuipers, O. P. Bistability, Epigenetics, and Bet-Hedging in Bacteria. Annu. Rev. Microbiol. 62, 193–210 (2008).

64. Roth Rosenberg, D., et al. Particle-associated and free-living bacterial communities in an oligotrophic sea are affected by different environmental factors. Environ. Microbiol. 23, 4295–4308 (2021).

65. Alcolombri, U. et al. Biogel scavenging slows the sinking of organic particles to the ocean depths. Nat. Commun. 16, 3290 (2025).

