## Supplementary Information for "TCA cycle entry point, growth variability and amino acid utilization in *Alteromonas macleodii* ATCC 27126"

SUPPLEMENTARY FIGURES:

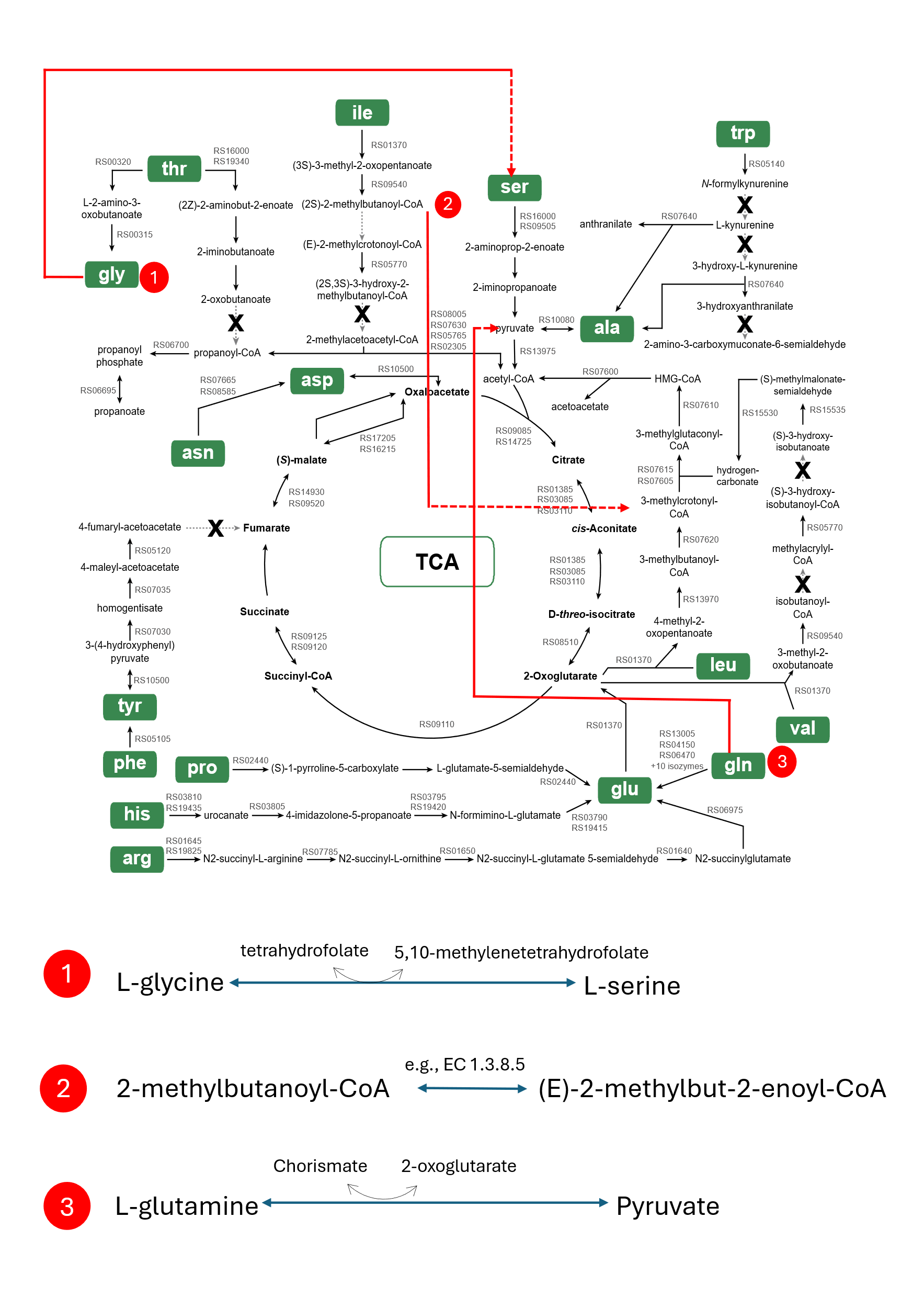

***Figure Supplementary 1: Amino acid degradation pathway maps (bioinformatic reconstructions). Reconstruction of catabolic pathways in A. macleodii 27126.***

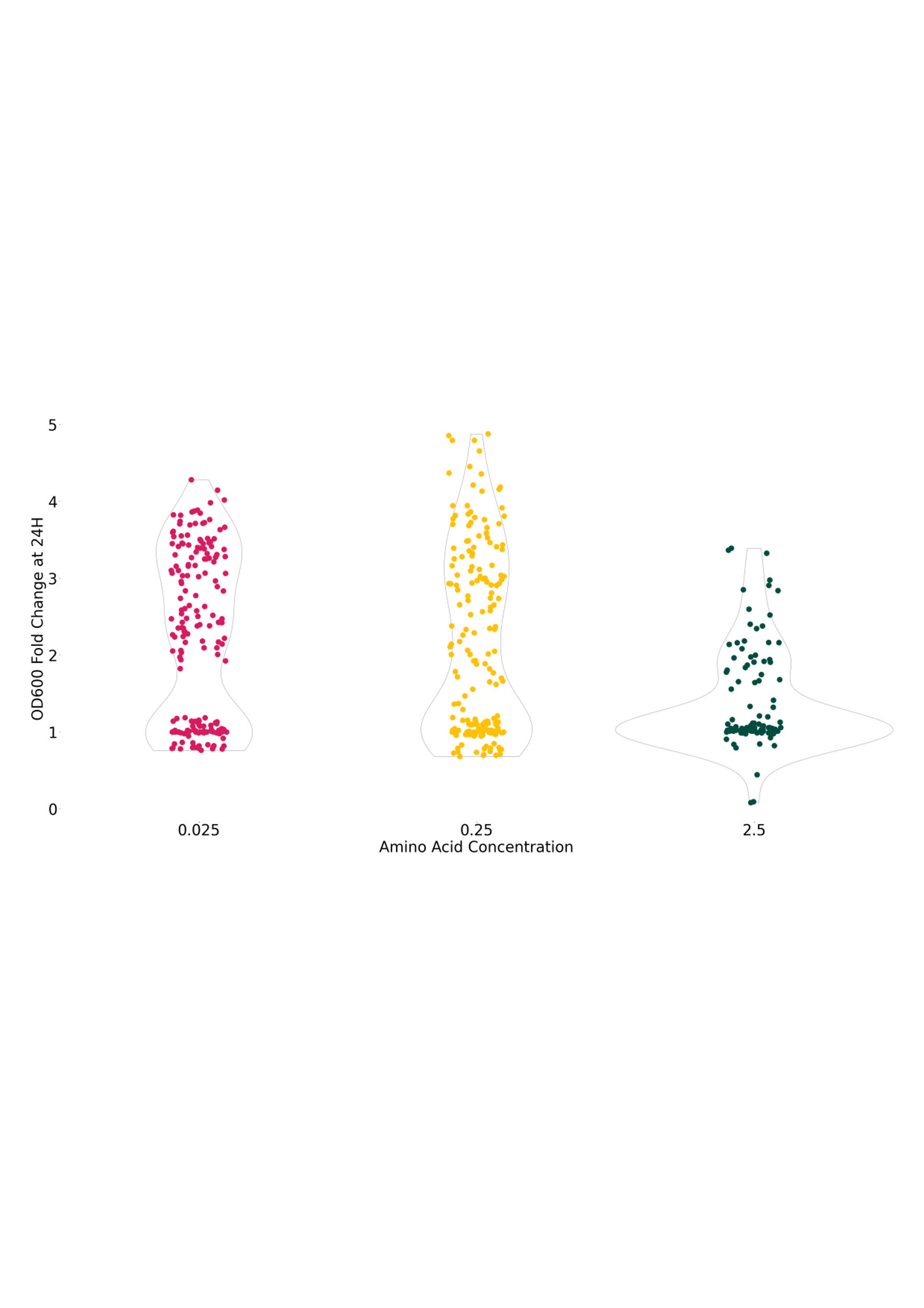

***Supplementary Figure 2: Experimental setup for amino acid concentrations and controls.*** *X axis shows the amino acid concentrations (% w/v). Y axis shows times fold change between 24-hour OD600 reading as opposed to the control samples.*

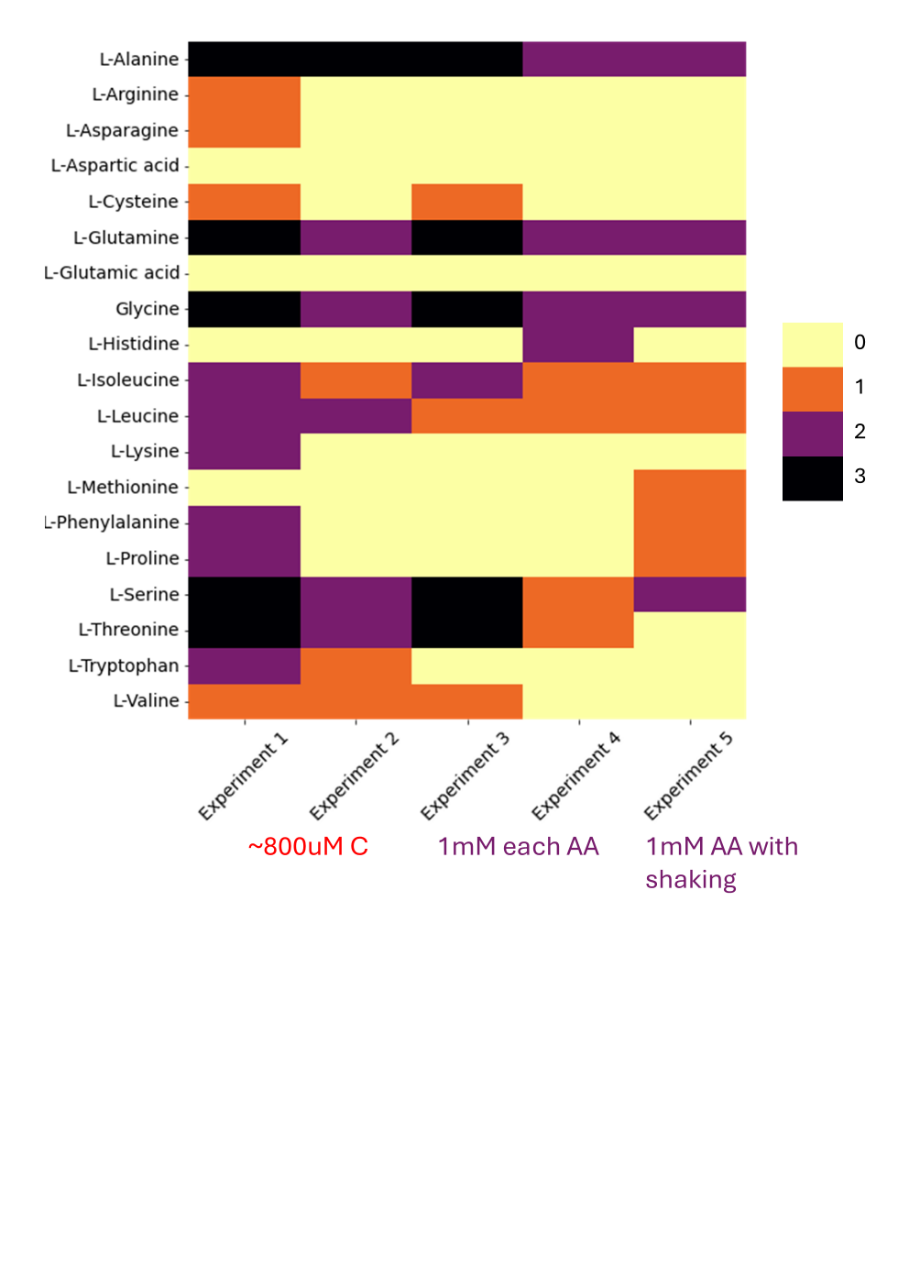

***Supplementary Figure 3: Replicate variability and probability of growth on individual amino acids.***

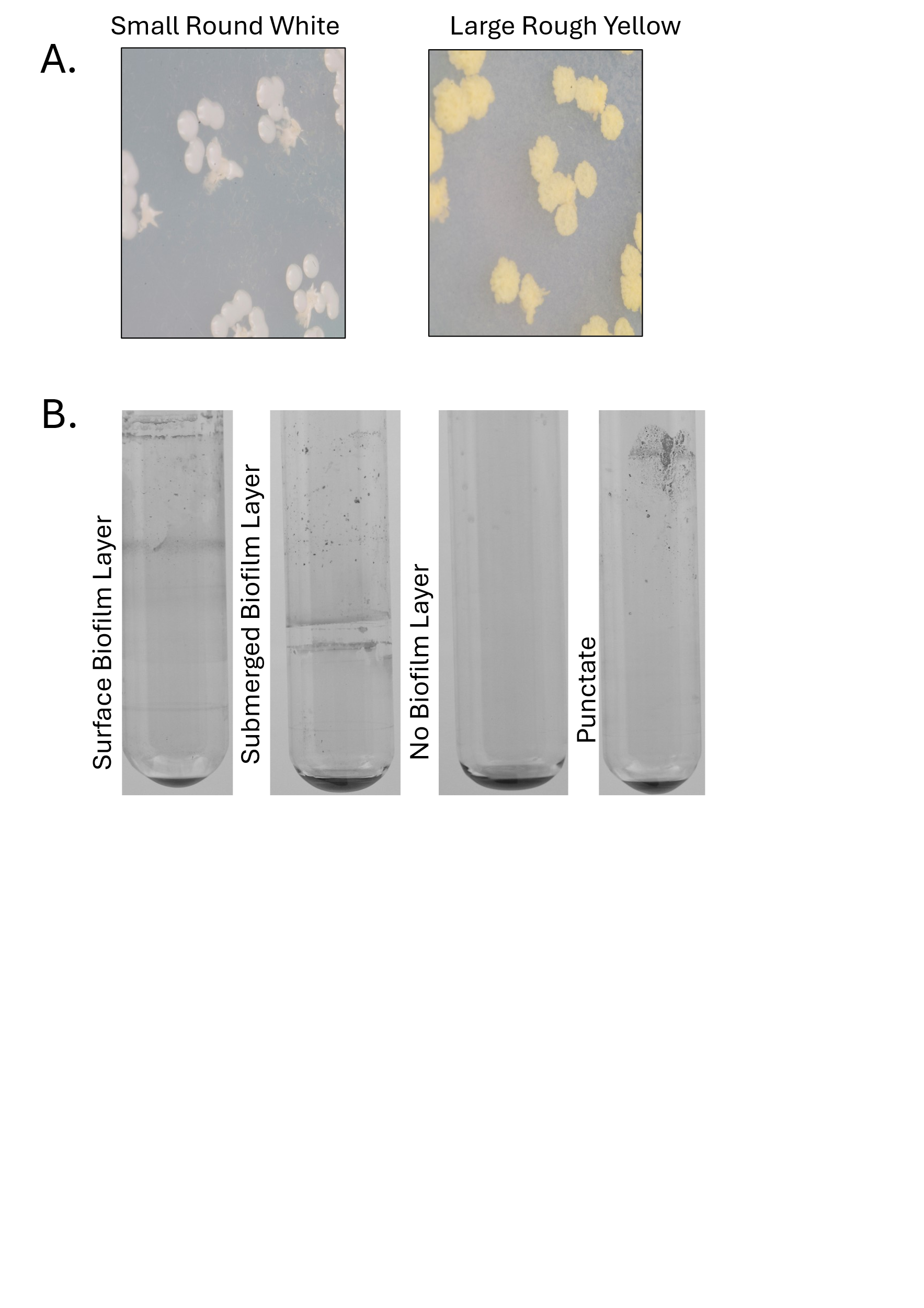

***Supplementary Figure 4: Photographs showing colony morphologies (LSW and SRY) and Biofilm formation patterns in test tube cultures.***
